# Measuring and modeling the dynamics of mitotic error correction

**DOI:** 10.1101/2024.01.10.574250

**Authors:** Gloria Ha, Paul Dieterle, Hao Shen, Ariel Amir, Daniel J. Needleman

## Abstract

Error correction is central to many biological systems and is critical for protein function and cell health. During mitosis, error correction is required for the faithful inheritance of genetic material. When functioning properly, the mitotic spindle segregates an equal number of chromosomes to daughter cells with high fidelity. Over the course of spindle assembly, many initially erroneous attachments between kinetochores and microtubules are fixed through the process of error correction. Despite the importance of chromosome segregation errors in cancer and other diseases, there is a lack of methods to characterize the dynamics of error correction and how it can go wrong. Here, we present an experimental method and analysis framework to quantify chromosome segregation error correction in human tissue culture cells with live cell confocal imaging, timed premature anaphase, and automated counting of kinetochores after cell division. We find that errors decrease exponentially over time during spindle assembly. A coarse-grained model, in which errors are corrected in a chromosome autonomous manner at a constant rate, can quantitatively explain both the measured error correction dynamics and the distribution of anaphase onset times. We further validated our model using perturbations that destabilized microtubules and changed the initial configuration of chromosomal attachments. Taken together, this work provides a quantitative framework for understanding the dynamics of mitotic error correction.

## Introduction

Error correction occurs in many biological systems ranging from DNA damage repair and protein synthesis to T-cell antigen binding(1–3). Error correction plays a critical role in protein function and cell health by promoting accurate protein interactions and fixing damaged and incorrect genetic material. Previous work in such systems has focused on understanding the tradeoffs between energy, speed, and accuracy in error correction through kinetic modeling and experimental measurements of the timing of molecular events (e.g., enzyme dwell times, cell division times) as well as baseline error rates(4–7). Measuring the dynamics of error correction in live cells is usually intractable due to the molecular scale and rapid speed of the processes involved. In this work, we tackle measurements and modeling of both the dynamics and resulting timing of mitotic error correction, a process which is essential for the faithful inheritance of genetic material.

The mitotic spindle is a bipolar structure, primarily composed of microtubules, that segregates an equal number of chromosomes to each daughter cell with remarkable fidelity. The spindle begins with many incorrect attachments between chromosomes and microtubules, which are corrected over time to eventually satisfy the spindle assembly checkpoint and allow the cell to proceed to segregate the chromosomes (8– 11). Chromosome missegregation is often attributed to defects in error correction (12–14). Despite the widespread importance of chromosome segregation errors, there is currently a lack of quantitative methods to characterize the dynamics of the error correction process and how it can go wrong.

Comprehensive measurement of chromosome segregation errors in human cells is challenging – past studies rely on identifying lagging chromosomes in anaphase or counting errors for one or two specific chromosomes(15, 16). While single cell sequencing(17, 18) and fixed cell labeling techniques(19) can determine which chromosomes missegregated, these static measurements do not provide information on the dynamics of the error correction process. Recent work has used high-resolution live cell imaging to track the movements of chromosomes from nuclear envelope breakdown to anaphase(20–22); but it is unclear to what extent chromosome alignment is an indicator of error status. To our knowledge, combining dynamic measurements during spindle assembly with accurate quantification of chromosome segregation errors has not been done, preventing an accurate understanding of how errors are corrected over time. Efforts have also been made to model the dynamics of spindle assembly and error correction(23–25), but these models are difficult to test experimentally due to the lack of detailed measurements of the many processes involved.

Here, we present a method to quantitively measure error correction dynamics in mitosis. We combine live-cell imaging of spindle assembly, high-resolution imaging of kinetochore counts, timed forced anaphase, and mathematical modeling. Our experimental results can be quantitatively described by a simple coarse-grained model, indicating that error correction is a chromosome autonomous process which occurs at a constant rate throughout the course of spindle assembly. Since anaphase commences when the last erroneously attached chromosome is corrected, solving for the distribution of anaphase times in this model is a slowest first passage problem, leading to a predicted non-trivial distribution that agrees with experimental observations. We perform additional experiments with chemical inhibitors and show that perturbations to microtubule stability and the initial configuration of the spindle can slow the rate of error correction. Taken together, this work provides a quantitative framework for understanding the dynamics of mitotic error correction.

## Results

### Kinetochore counting as a measure of chromosome segregation errors

We sought to develop a live-cell imaging method to measure chromosome segregation errors by counting the number of kinetochores in each daughter cell after division. We used the chromosomally stable diploid human tissue culture cell line hTERT-RPE-1 that we constructed to stably express sfGFP::CENP-A, allowing for clear imaging of kinetochores during mitosis and after division. We also inserted mCherry::alpha-tubulin in the same cell line to visualize spindle microtubules during mitosis (Fig 1A). We stained the cells with SPY650-DNA to identify the timing of chromosome condensation and nuclear envelope breakdown. After cell division, we took 0.5 µm spaced, high laser intensity z-stacks of the divided cells’ kinetochores (Fig 1B).

**Figure 1.**
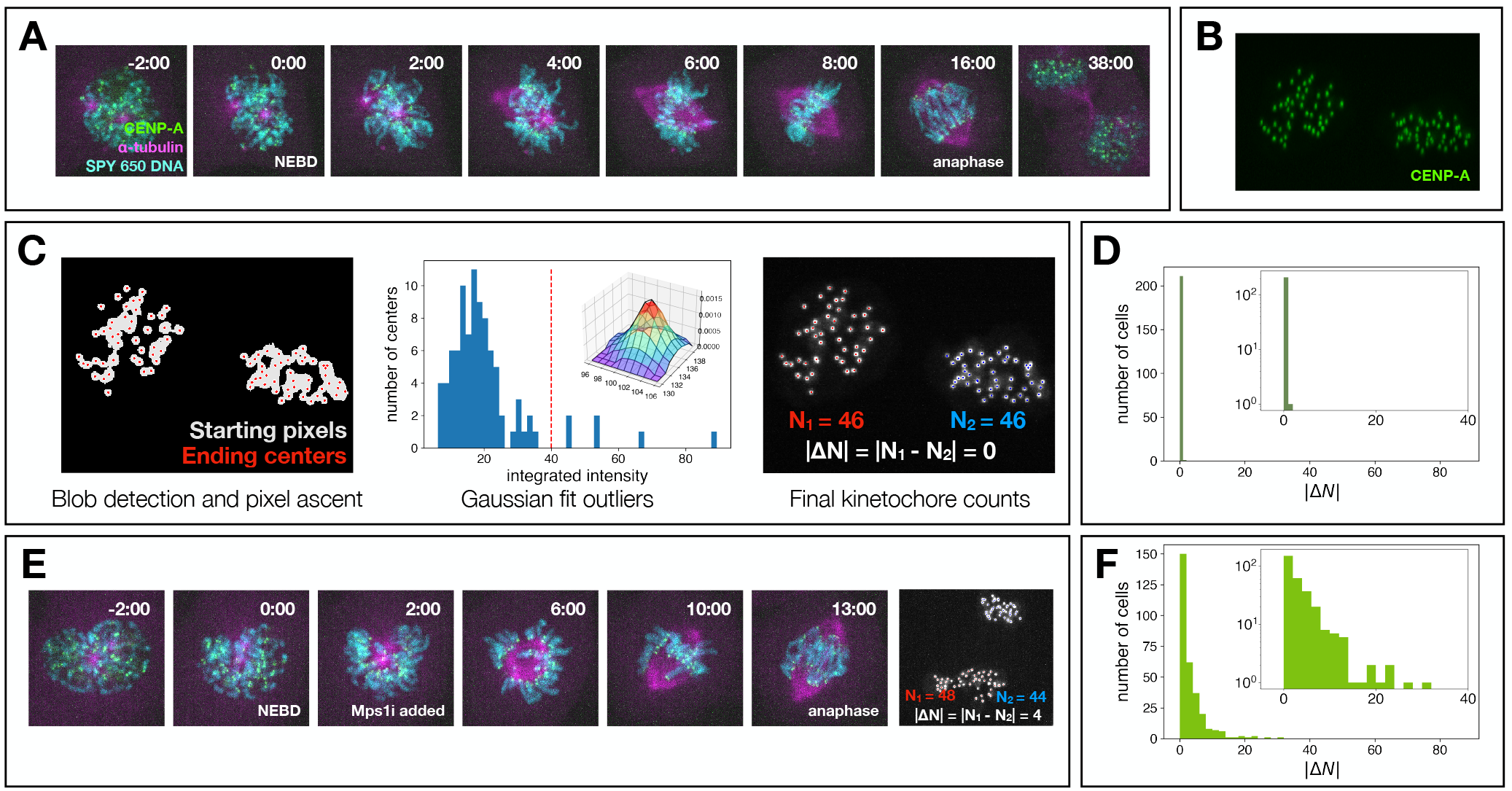
Kinetochore counting as a measure of chromosome segregation error. **(A)** Mitosis time-lapse of RPE-1 cell expressing sfGFP-CENP-A and mCherry-α-tubulin and dyed with SPY 650 DNA (time 0 is end of nuclear envelope breakdown). **(B)** Maximum intensity projection of z-stack of CENP-A (kinetochore marker), same cell as (A). **(C)** Image analysis pipeline to count kinetochores: potential kinetochores are identified and kinetochore counts are finalized by manual investigation of Gaussian fit integrated intensity outliers (kinetochore clusters), yielding kinetochore difference metric |ΔN|. **(D)** Histogram of |ΔN| values for unperturbed control cells (n=212 cells), with the inset showing the same data plotted on a log-linear scale. **(E)** Mitosis timelapse of RPE-1 cells with prematurely forced anaphase (addition of Mps1 inhibitor AZ-3146) and resulting kinetochore counts. **(F)** Histogram of |ΔN| values for cells forced into anaphase after NEBD (n=299 cells), with the inset showing the same data plotted on a log-linear scale.

To quantify the number of kinetochores in the divided cells, we developed an image analysis algorithm to locate kinetochores in the 3D z-stacks (Fig 1C). We first used a Gaussian fit of the pixel intensities in the green channel (sfGFP::CENP-A) to mask the cytoplasm area from the background. We then used median thresholding of the intensity distribution of the pixels in the cytoplasmic mask to threshold the green channel for kinetochore-containing pixels. We used a gradient ascent method where “walkers” were initially placed at each pixel in the kinetochore threshold and the walkers subsequently stepped in 3D to the brightest neighboring pixel until their positions stabilized at local intensity maxima. Final walker positions that satisfied thresholds for pixel intensity and number of walkers were taken as the candidate kinetochore centers. We then used k-means clustering with k=2 to assign candidate centers to the two daughter cells, resulting in preliminary kinetochore counts for each of the daughter cells. We performed 3D Gaussian fits on all the candidate kinetochores and plotted the distribution of their integrated intensities. We manually investigated outliers in integrated intensity by scrolling through the original green channel z-stack and checking whether any of the candidate centers that had high integrated intensities were clusters of two or more kinetochores. This procedure allowed us to measure the number of kinetochores in the two daughter cells (*N*_1_, *N*_2_). We repeated this measurement in 214 RPE-1 cells and calculated the total number of kinetochores (*N*_*1*_ + *N*_*2*_) per daughter cell pair, with all but 2 cells giving 92 kinetochores. Such a high rate of euploidy is expected for RPE-1 cells(15), which argues that the developed imaging and analysis provides a reliable measure of the true number of kinetochores.

We next investigated the kinetochore count difference |*ΔN*| = *N*_*1*_ -*N*_*2*_ between daughter cells with 92 kinetochores. Only one of 212 RPE-1 cells had a nonzero kinetochore count difference (Fig 1D). If we naively assume that cells with |*ΔN*| = 0 contain no segregation errors, and that the one cell with |*ΔN*| = 2 had one segregation error, that gives a lower bound error rate of one error in 212 divisions. This estimate is consistent with previous fluorescence in-situ hybridization measurements in RPE-1 cells indicating an error rate of approximately one error in 100 divisions(15, 17).

We next investigated how prematurely entering anaphase impacts chromosome segregation errors. We first treated unsynchronized, mitotic cells with the spindle assembly checkpoint inhibitor AZ-3146, which inhibits Mps1 (26). Previous studies have shown that inhibiting Mps1 in mitotic cells before anaphase onset leads to errors in chromosome segregation(27). When we inhibited Mps1 after the end of nuclear envelope breakdown (NEBD), cells went into anaphase 6.2±0.2 minutes after Mps1 inhibitor addition on average, some with clearly uncentered and/or lagging chromosomes (Fig 1E). Out of 299 cells where we forced anaphase after NEBD, 149 cells (49.8%) had nonzero |*ΔN*| (Fig 1F), compared to 0.5% of cells in the unperturbed case above (Fig 1D). Thus, prematurely forcing cells to enter anaphase before there is adequate time to correct errors results in a highly elevated frequency of chromosome segregation errors.

### Time course forced anaphase measurements and coarse-grained model of error correction dynamics

We next sought to use forced anaphase to measure error correction dynamics during normal spindle assembly. Since we recorded movies of all cells from before nuclear envelope breakdown (NEBD), we were able to determine the time elapsed between NEBD and Mps1i inhibitor addition for each cell (Fig 2A), as well as the time between Mps1 inhibitor addition and anaphase onset. The time between Mps1 inhibitor addition and anaphase onset was longer for cells that were forced into anaphase shortly after NEBD (*t*_ana_=8.8±0.3 minutes for cells forced between 0 and 5 minutes post NEBD) than cells that were forced later (*t*_ana_=3.8±0.2 minutes for cells forced between 10 and 15 minutes post NEBD), which may be due to cellular processes that must occur before anaphase can begin setting a minimum mitotic duration time. If Mps1 inhibitor addition “freezes” error correction, as our subsequent results indicate (see below), then error correction only occurs in the time interval between NEBD and Mps1 inhibitor addition. We thus considered cells that were forced into anaphase via Mps1i addition between 0 and 16 minutes after NEBD, since unforced RPE-1 cells spontaneously enter anaphase 17.6±0.3 minutes after NEBD, and grouped cells into 4 minute time intervals and made histograms of their |*ΔN*| values. The |*ΔN*| histograms were very broad for early timepoints, when the cell hadn’t had time to correct many attachments, and narrower for later timepoints, with increasing fractions of cells with |*ΔN*| = 0 (Fig 2B). This result is consistent with errors being corrected over time spent in spindle assembly.

**Figure 2.**
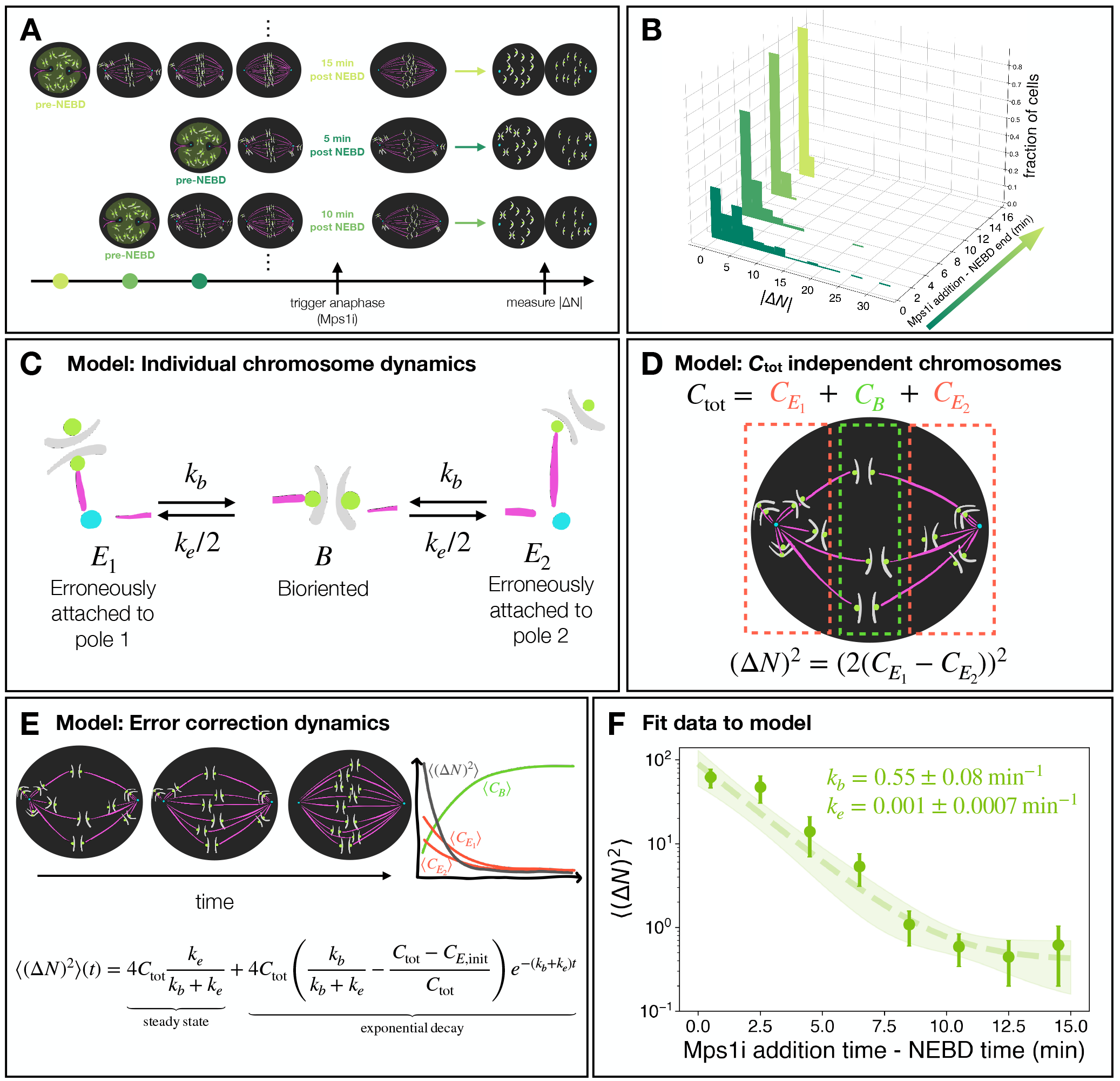
Coarse-grained modeling of kinetochore count dynamics to extract error correction rates from data. **(A)** Diagram of forced anaphase assay on unsynchronized cells. **(B)** Histograms of |ΔN| for each forced anaphase time (n=[102,86,71,40] cells for t<[4,8,12,16] minutes). **(C)** Three state diagram for chromosomes independently correcting and becoming erroneous. **(D)** Connection of state model to experimentally measured kinetochore count difference metric. **(E)** Schematic of time dynamics of error correction and model solution. Steady state of the mean squared kinetochore count difference is given by the ratio of correction rates and the exponential decay depends on the correction rate. **(F)** Plot of experimental <(ΔN)^2^>(t) binned in 2 minute intervals (error bars are standard error of the mean) and model fit (shaded area represents two standard deviations above and below the best fit curve).

We next developed a coarse-grained model of the dynamics of error correction to aid in interpreting our experimental results. In this model, each cell contains a total of *C*_tot_ chromosomes, each of which can be in three states: bioriented (*B*), erroneously attached to pole 1 (*E*_1_), or erroneously attached to pole 2 (*E*_2_). A chromosome in state *B* is defined as one in which the two sister chromatids will correctly segregate into the two daughter cells upon entry into anaphase, while chromosomes entering anaphase in states *E*_1_ or *E*_2_ will result in both chromatids in the same daughter cell. We used the simplest assumption for error correction dynamics: that the chromosomes are independent of each other and that chromosomes in states *E*_1_ and *E*_2_ are corrected with constant rate *k*_*b*_, and that chromosomes in state *B* become erroneously attached with constant rate *k*_*e*_ (Fig 2C).

In this model, the total number of chromosomes in states *B, E*_1_, and *E*_2_ at time *t* are *C*_*B*_(*t*), 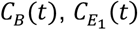, and 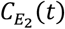 respectively (Fig 2D). If the cell were forced into anaphase at time *t*, it would have a difference in the number of kinetochores between daughter cells of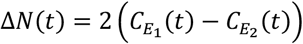, where the factor of two results from there being two kinetochores per chromosome. Using a matrix equation approach, and assuming equal probabilities for an initially erroneously attached chromosome of being in state *E*_1_ and *E*_2_, we solved for the dynamics of error correction in this model (Eq. 1, SI Appendix; Fig. 2E), and found

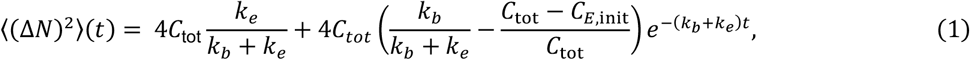

where *C*_*E, init*_ is the average initial number of erroneously attached chromosomes at the beginning of spindle assembly. Thus, ⟨(Δ*N*)^2^⟩(*t*) decays exponentially over time, with a time scale set by the sum of the biorientation rate *k*_*b*_ and the error rate *k*_*e*_, approaching a steady-state value given by the ratio of *k*_*b*_ to *k*_*e*_. This predicted form of the error correction dynamics (Eq. 1) does an excellent job of describing the experimentally-measured time course of ⟨(Δ*N*)^2^⟩ (Fig. 2F), with a fit giving *k*_*b*_ = 0.55 ± 0.08 min^-1^ and *k*_*e*_ = 0.001 ± 0.0007 min^-1^. Thus, the rate of error correction is much higher than the rate of forming new errors (*k*_*b*_ ≫ *k*_*e*_) in unperturbed RPE-1 cells, which is consistent with RPE-1 being a chromosomally stable and euploid cell line.

Since Mps1 plays a role in error correction outside of spindle assembly checkpoint timing(28, 29), we sought to test whether the elevated errors observed upon Mps1i addition were solely due to early entry into anaphase or, alternatively, if errors were further elevated by interfering with other functions of Mps1. We thus investigated an alternative method of bypassing the spindle assembly checkpoint. We targeted Mad2, whose localization to erroneously attached kinetochores prevents anaphase onset(30, 31). We knocked down Mad2 via RNA interference treatment, which has previously been shown to cause premature anaphase and chromosome segregation errors in cells(32, 33). Mad2 RNAi cells entered anaphase 9.7±0.3 minutes after nuclear envelope breakdown (Fig S1Aii) and the divided cells had a ⟨(Δ*N*)^2^⟩ of 14.3±0.4 (Fig S1Ai). The ⟨(Δ*N*)^2^⟩ for Mad2 RNAi cells is higher than the ⟨(Δ*N*)^2^⟩ for the cells for which Mps1i was added around the same time (⟨(Δ*N*)^2^⟩(*t* = 10.5 minutes) = 0.6 ± 0.2, p < 0.01, Mann WhitneyU). Shifting the Mad2 RNAi ⟨(Δ*N*)^2^⟩ back by 5 minutes makes the Mad2 RNAi overlap with the Mps1 inhibitor data (Fig S1Aiii, p<0.83, Mann WhitneyU), which is not unreasonable given the 6.2±0.2 minute lag time we observed between Mps1i addition and anaphase onset. The agreement between the Mad2 knockdown data with the Mps1i data indicates that Mps1i does not unduly inflate errors.

### Slowest first passage times of the coarse-grained model predicts the distribution of anaphase times and agrees with error correction data

We next sought alternative means to test if the simple coarse-grained model correctly describes the dynamics of error correction. When the spindle assembly checkpoint is functional, anaphase only begins after the last chromosome is properly attached to the spindle (at time *t*_*f*_), which, after an offset time (*t*_offset_), results in the initiating of chromosome segregation which can be visualized with microscopy (at time *t*_*f*_ + *t*_offset_) (Fig. 3A). In the coarse-grained model (Fig 2C), *t*_*f*_ corresponds to the first time at which all of the *C*_tot_ chromosomes in the spindle are in the correctly attached state *B*. Since our measurements show that the rate at which erroneous attachments are corrected is much faster that the rate at which correct attachments become erroneous (i.e. *k*_*b*_ ≫ *k*_*e*_), we neglect transitions from state *B* to the *E* states in a first approximation. Thus, the time at which the last chromosome enters state *B* becomes a slowest first passage time problem, which can be shown to follow a Gumbel distribution(34, 35) (SI Appendix). If there are initially *C*_*E, init*_ erroneously attached chromosomes, then the distribution of such slowest first passage times *t*_*f*_ is

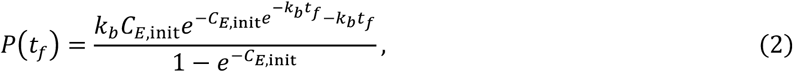

which leads to the coarse-grained model’s prediction for the distribution of observed anaphase times (*t*_*f*_ + *t*_offset_). Moments of the Gumbel distribution can be analytically calculated, revealing that while the mean anaphase time depends on all 3 parameters (*k*_b_, *C*_*E*, init_, *t*_offset_), the variance is solely determined by the error correction rate *k*_*b*_: with a variance in anaphase times of 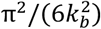 (SI Appendix). Thus, fitting anaphase times to the Gumbel distribution can well constrain *k*_*b*_.

**Figure 3.**
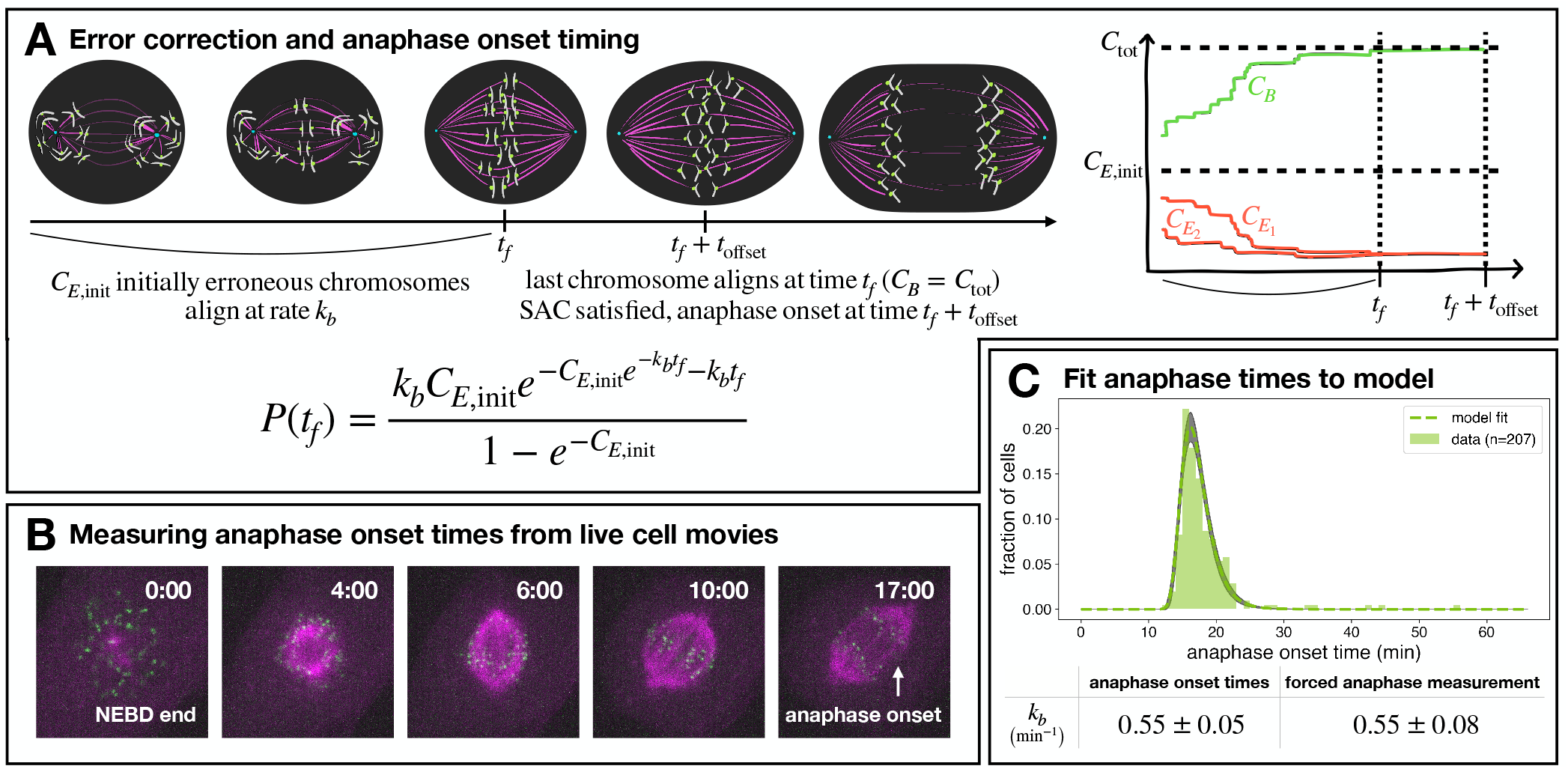
Anaphase timing model agrees with error correction results. **(A)** Schematic relating error correction, anaphase onset time, and coarse-grained model from Fig. 2C. **(B)** Timelapse images of RPE-1 cells undergoing spindle assembly with anaphase onset time marked. **(C)** Anaphase onset times fit to an analytical distribution of the slowest first passage time (n=207).

We next measured anaphase onset times from live cell movies of RPE-1 spindle assembly, defining anaphase onset time as the time from when nuclear envelope breakdown ends to when the kinetochores start separating (Fig. 3B). The experimentally measured distribution of anaphase onset times is well fit by Eq (2), with a best fit giving *k*_*b*_ =0.55±0.05 min^-1^ (Figure 3C). This *k*_*b*_, determined from fitting the distribution of anaphase onset times, is within error the same as the *k*_*b*_ =0.55±0.08 min^-1^ determined from fitting from the kinetochore count error data (Fig. 3C, p < 0.96, Student’s t-test). Simultaneously fitting the anaphase time and kinetochore count data also yielded *k*_*b*_ =0.55±0.02 min^-1^ (Table S1). That these two orthogonal methods of measuring *k*_*b*_ are consistent with each other provides strong evidence for the validity of the simple coarse-grained model of error correction dynamics. The simultaneous fit also gave *C*_*E*, init_ = 26 ± 3 chromosomes, which is consistent with each kinetochore initially randomly and independently associating with either pole: in such a scenario there would be a 50% chance of both kinetochores from each chromosome being bioriented, and a 50% chance of both kinetochores attaching to the same pole, giving an expected *C*_*E*,init_ of 0.5 × 46 = 23 chromosomes (quite similar to the measured value).

### Error correction rate depends on microtubule stability and spindle assembly pathway

We next investigated how both anaphase onset timing and error correction dynamics change in response to molecular perturbations. Since the slowest first passage time model for anaphase timing is only valid when the spindle assembly checkpoint is functional, we aimed to modulate the rate of error correction without perturbing the checkpoint. We first used UMK57, a small molecule potentiator of the kinesin-13 motor MCAK, which induces increased microtubule detachment from kinetochores(36, 37). When imaging cells undergoing spindle assembly in 1µM UMK57, we saw that cells took longer to enter anaphase (Fig 4A). This is consistent with previous work in other cell lines that show that saturating doses of UMK57 lengthen mitotic duration(36). We measured kinetochore count differences in UMK57 cells that were not forced into anaphase and saw that 3 out of 110 cells (2.7%) had a nonzero kinetochore count difference, indicating a low baseline rate of errors in the presence of UMK57 (Fig 4B).

**Figure 4.**
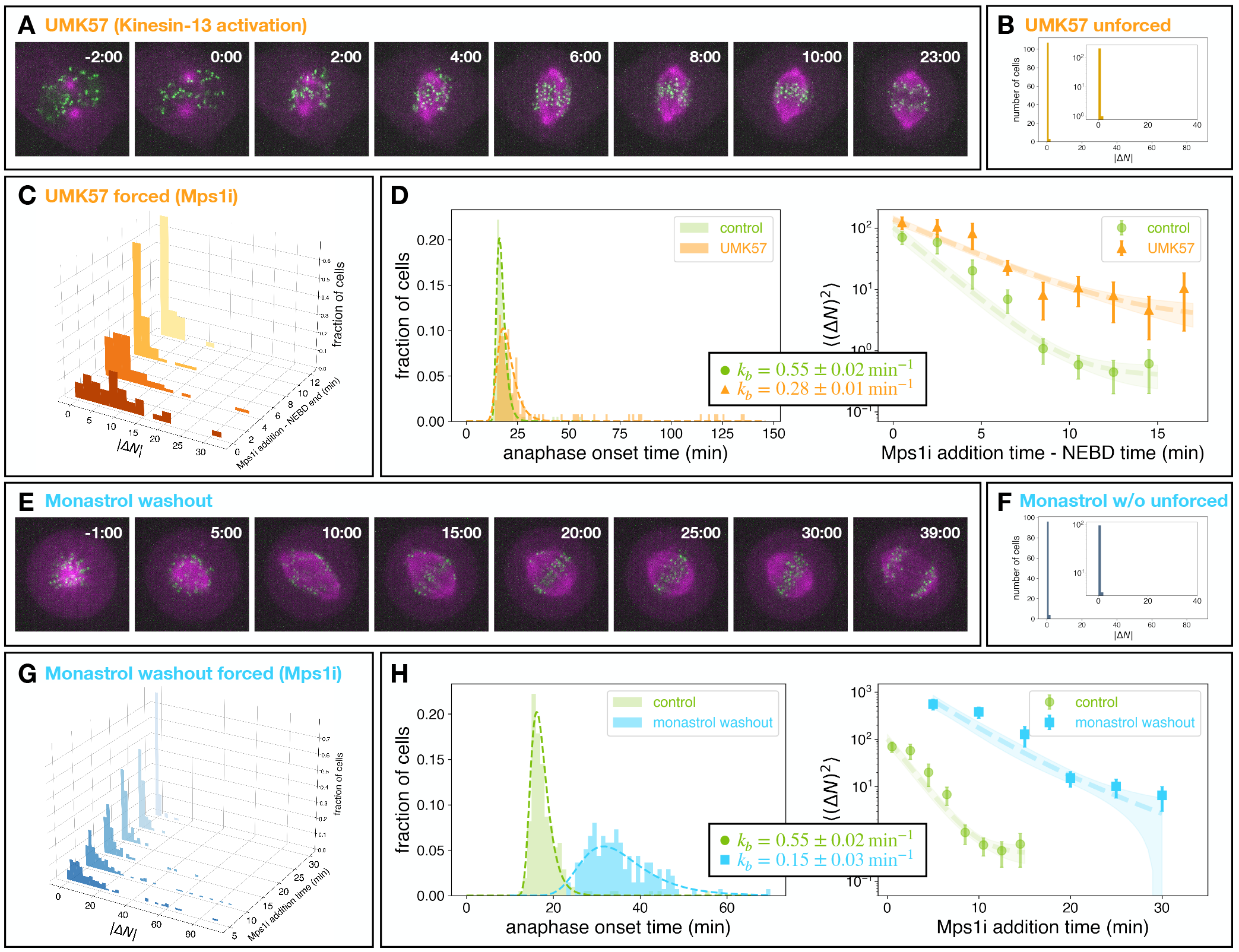
Perturbing microtubule stability, initial configuration affect error correction. **(A)** Time-lapse of unsynchronized cells subject to 1uM UMK57. **(B)** Histogram of |ΔN| values for unforced UMK57 cells (n=110 cells), with the insert showing the same data plotted on a log-linear scale. **(C)** Histograms of |ΔN| for each UMK57 forced anaphase time (n=[35,59,60,38] cells for t<[4,8,12,16] minutes). **(D)** Anaphase time histogram (n=128) and kinetochore count error correction curve for UMK57 (orange) vs control (green). *k*_*b*_ fit is from simultaneous fit to both anaphase time and kinetochore count data. **(E)** Time-lapse of cell undergoing monastrol washout. **(F)** Histogram of |ΔN| values for unforced monastrol washout cells (n=100 cells), with the inset showing the same data plotted on a log-linear scale. **(G)** Histograms of |ΔN| for each monastrol washout forced anaphase time (n=[99, 138, 70, 53, 61, 69] cells for t=[5, 10, 15, 20, 25, 30] minutes). **(H)** Anaphase time histogram (n=137) and kinetochore count error correction curve for monastrol washout (blue) vs. control (green). *k*_*b*_ fit is from simultaneous fit to both anaphase time and kinetochore count data.

We repeated the forced anaphase experiment (Fig 4C) and measured anaphase times in unforced UMK57 cells (Fig 4D, left). We used the predictions of the coarse-grained model (Eqs. 1 and 2) to simultaneously fit both the distribution of anaphase times and the error correction time course of ⟨(Δ*N*)^2^⟩(*t*), providing an excellent fit to the two data sets (Fig 4D, orange dashed lines). The resulting rate of error correction in cells exposed to UMK57 was *k*_*b*_ =0.28±0.01 min^-1^, which is substantially smaller than controls (for unperturbed RPE-1 cells, simultaneous fitting of the distribution of anaphase times and the error correction time course gives *k*_*b*_ =0.55±0.02 min^-1^, p<0.0001, Student’s t-test) (Table S1). Individually fitting the UMK57 datasets yielded similar results to the simultaneous fit, with *k*_*b*_ =0.32±0.01 min^-1^ for the forced anaphase experiment and *k*_*b*_ =0.27±0.01 min^-1^ for the unforced anaphase times. Thus, increasing microtubule detachment from kinetochores with UMK57 slows down the rate of error correction, leading to prolonged anaphase times.

In order to cross-check our Mps1i results for UMK57-treated cells with an alternative spindle assembly checkpoint-bypassing method, we characterized Mad2 RNAi cells treated with UMK57. The average anaphase onset time in UMK57-treated cells decreased from 30±2 minutes without Mad2 RNAi to 11.1±0.4 minutes with Mad2 RNAi (Fig S1Bii). As in control RPE-1 cells, shifting the ⟨(Δ*N*)^2^⟩ for UMK57-treated Mad2 RNAi back 5 minutes caused it to overlap with the ⟨(Δ*N*)^2^⟩ from UMK57-treated cells forced into anaphase via Mps1 inhibitor (Fig S1Biii, p<0.15, Mann WhitneyU), once again indicating that the results with Mps1i were comparable to those with Mad2 knockdown.

Arresting cells in mitosis with the Eg5 inhibitor monastrol causes spindles to become monopolar and induces erroneous attachments which have been used to study error correction(38–40). We next sought to determine how inducing such an error prone state impacts the dynamics of error correction. We arrested cells using a 2-hour incubation in the Eg5 inhibitor monastrol. After an on-scope washout from monastrol incubation, spindles swiftly bipolarized and cells divided with a mean time of 35±1 minutes (Fig 4E). The resulting kinetochore count differences were low – 4% of cells had a nonzero |*ΔN*|, which is consistent with error rates in RPE-1 cells after monastrol washout previously measured by single cell sequencing and fluorescence in-situ hybridization(15, 17) (Fig 4F).

To test how forcing premature anaphase affected monastrol-arrested cells undergoing spindle bipolarization, we again used the Mps1 inhibitor AZ-3146. We forced anaphase at defined points in time after monastrol washout, from 5 to 30 minutes. The histograms of |ΔN| values were very broad for early timepoints, when the cell hadn’t had time to correct many attachments, and narrower for later timepoints, with increasing number of cells with |ΔN| = 0 as time between monastrol washout and Mps1 inhibitor addition increased (Fig 4G). This result is consistent with errors being corrected over time spent in spindle bipolarization. The initial ⟨(Δ*N*)^2^⟩ (5 minutes after monastrol washout) was much larger in magnitude than that of either the control or the UMK57 (MCAK potentiator) condition. If the initially erroneous attachments are equally likely to be associated with either pole, then the maximum possible initial ⟨(Δ*N*)^2^⟩ is 184 (SI Appendix), which is far less than the experimental measured initial ⟨(Δ*N*)^2^⟩ in the monastrol washout (558±142, p<0.0005, bootstrapping). We thus expanded the coarse-grained model to account for a statistical asymmetry in the initial erroneous attachments (i.e. that cells can contain a “stronger” pole that initially erroneous attachments are more likely to be associated with), and also extended the anaphase timing predictions to account for a finite backwards rate *k*_*e*_ (SI Appendix).

We used the predictions of the coarse-grained model with initially statistically asymmetric attachments and finite *k*_*e*_ to simultaneously fit both the distribution of anaphase times and the error correction time course of ⟨(Δ*N*)^2^⟩(*t*), providing an excellent fit to the two data sets (Fig 4H, blue dashed lines). The resulting rate of error correction in cells after monastrol washout was *k*_*b*_=0.15±0.03 min^-1^, which is substantially smaller than the control (for unperturbed RPE-1 cells *k*_*b*_=0.55±0.02 min^-1^, p<0.0001, Student’s t-test; fitting with the expanded coarse-grained model, allowing for initial asymmetry and finite *k*_*e*_ in anaphase timing, did not substantially change the parameters for control and UMK57, Table S1). Fitting the two monastrol washout datasets separately yielded similar results to the simultaneous fit, with *k*_*b*_=0.13±0.02 min^-1^ for the forced anaphase experiment and *k*_*b*_=0.18±0.01 min^-1^ for the unforced anaphase times. Thus, arresting cells as monopolar spindles slows down the rate of error correction, leading to prolonged anaphase times.

As a control for Mps1i, we once again compared these monastrol washout results to those from washing out monastrol in cells treated with Mad2 siRNA. Mad2 knockdown via RNAi decreased the mean anaphase onset times from 35±1 minutes without RNAi to 20±1 minutes with RNAi (Fig S1Cii). Unlike control Mad2 RNAi cells and UMK57-treated Mad2 RNAi cells, anaphase onset times for monastrol washout cells with Mad2 RNAi spanned a wide range (9 to 43 minutes). We thus were able to calculate the ⟨(Δ*N*)^2^⟩ for cells with similar anaphase onset times (4 minute bins) for monastrol washout cells with Mad2 RNAi, and found that ⟨(Δ*N*)^2^⟩ decreased over time in the Mad2 RNAi data in a manner that was qualitatively similar to the Mps1i forced anaphase data (Fig S1Ciii), again indicating that the Mps1i results were consistent with bypassing the spindle assembly checkpoint with Mad2 knockdown.

Taken together, our results demonstrate that a simple coarse-grained model in which erroneous attachments are corrected in a chromosomes autonomous manner at a constant rate can recapitulate both the experimentally observed error correction rates and anaphase onset times, and can be used to study how these change in response to molecular perturbation.

## Discussion

In this study, we developed a method to quantitatively measure and model mitotic error correction dynamics in human tissue culture cells by combining kinetochore count and anaphase timing measurements with a coarse-grained model of error correction. Previous work on measuring errors and error correction has focused either on endpoint chromosome segregation error measurements or on tracking chromosome positions over time(15, 17, 20). By combining live cell imaging of mitotic progression with our timed segregation error measurement, we quantitatively measure the dynamics of error correction.

Our method entails forcing cells into anaphase at different points after nuclear envelope breakdown and observing how the kinetochore count difference changes with time spent in spindle assembly. We used the Mps1 inhibitor AZ-3146 to trigger premature anaphase. Since Mps1 has a role in error correction in addition to the spindle assembly checkpoint (26), there is the potential concern that Mps1 inhibition might enhance chromosome segregation errors beyond merely inducing early entry into anaphase. However, the observed reduction in errors with time spent in bipolarization/spindle assembly must be due to processes endogenous to the cells since all cells forced into anaphase are exposed to Mps1 inhibition, just applied at different times relative to NEBD. Additionally, comparison experiments measuring errors under Mad2 RNAi in all three conditions tested (control, UMK57, and monastrol washout), showed that the results obtained in the Mps1i experiments were consistent with bypassing the spindle assembly checkpoint through an alternative method. Furthermore, the error correction rate we measured from forcing cells into anaphase are the same as the error correction rate we inferred from the distribution of anaphase onset times (which involves no Mps1 inhibition), further arguing that Mps1 inhibition does not unduly inflate errors. Given the agreement of the error correction rate measured from the forced anaphase data with both the anaphase time distribution and Mad2 RNAi data, we believe that Mps1i “freezes” the error correction process soon after drug addition, independent of the varying lag times between drug addition and anaphase onset.

We found that incubation with UMK57 (MCAK activator) and washout from monastrol arrest both decreased the rate of error correction. While UMK57 in low doses has been shown to shorten mitotic duration in chromosomally unstable cells(36, 41), we observed that a high dose of UMK57 slowed down mitotic duration in chromosomally stable RPE-1 cells, presumably due to an increase in kinetochore microtubule instability leading to an impaired ability to correct uncentered chromosomes. We found that the initial average ⟨(Δ*N*)^2^⟩ for a monastrol washout was an order of magnitude higher than that of either unperturbed cells or cells exposed to UMK57, while the error correction rate was three-fold slower. Monopolar cells are thought of as an “error-prone” initial condition in part due to having a high proportion of monotelic and syntelic erroneous attachments (39), so the higher initial number of erroneous attachments in the monastrol washout was expected. It is less clear why the rate of error correction is slower after monastrol washout. It is possible that the type of errors that occur in monopolar spindle are different from, and corrected more slowly than, the type of errors that occur in unperturbed spindle assembly. Alternatively, prolonged metaphase arrest in a monopolar state may lead to abnormal stabilization of kinetochore fibers after monastrol washout due to perturbed cell cycle timing (42–44), leading to a slower error correction rate.

The quantitative agreement between our simple coarse-grained model and our experimental data indicates that error correction is a chromosome autonomous process that occurs at a constant rate over the course of spindle assembly. When the spindle assembly checkpoint is properly functioning, anaphase commences when the last erroneous chromosome becomes corrected, so solving for the distribution of anaphase times in this model amounts to calculating the maximum of many independent variables (i.e. the slowest chromosome to correct). This is a well-understood mathematical problem, and in the case where the independent variables are not bounded and their tails decay sufficiently fast, the solution is the Gumbel distribution (35, 45). The Gumbel distribution has been shown to describe a range of other biological phenomena, including menopause timing, bacteria lag time distributions, and actin cable length distributions in yeast(34, 46, 47). Despite the self-evident large differences between these different phenomena, their quantitative similarities highlight the common principles that dictate their behaviors.

The coarse-grained model developed here can quantitatively explain the experimentally observed error correction rates and anaphase onset times. By fitting one or both of those data sets, it is possible to extract the error correction rate, *k*_*b*_. Recent studies have shown that certain chromosomes are more likely to missegregate depending on their size and initial position in the nucleus (17, 20, 48, 49), which suggests that error correction rates may differ between chromosomes. If correct, this would imply that the error correction rate we measure is really an “effective” rate that is averaged over all chromosomes. It would be an interesting future direction to expand the current measurements and modeling to further explore these chromosome-specific differences. Additionally, the error correction rate, *k*_*b*_, presumably depends on many biophysical factors such as kinetochore microtubule stability, molecular motors, and Aurora B activity (43, 50–53). Determining how different molecular perturbations quantitively impact these biophysical factors and *k*_*b*_ will provide a means to test detailed models of the mechanistic basis of error correction (24, 54).

Previous experimental work on error correction in other biological systems has largely focused on measuring average error rates and timing(5, 6), but measurement of the dynamics of error correction in live cells has been largely inaccessible. In this work, we measured the time dynamics of mitotic error correction, in addition to error rates and distribution of anaphase timing. We found that despite the complex network of protein interactions that drive mitotic error correction, the overall dynamics of errors and anaphase timing can be explained by a minimal model in which errors are corrected independently at a constant rate. We were able to validate our model of error correction dynamics by measuring dynamics in two ways. A similar strategy could be informative in other contexts – combining this work with quantitative measurements of the dynamics of other error correction processes could lead to general insights into the principles that govern the dynamics of error correction in biology.

## Materials and Methods

### Cell lines

hTERT RPE-1 cell lines were maintained in Dulbecco’s modified Eagle’s medium GlutaMAX (DMEM GlutaMAX, Thermo Fisher) supplemented with 10% Fetal Bovine Serum (FBS, Thermo Fisher) and 50 IU ml^-1^ penicillin and 50 ug ml^-1^ streptomycin (Thermo Fisher) at 37°C in a humidified atmosphere with 5% CO_2_. Cells were regularly validated as mycoplasma free by a PCR-based mycoplasma detection kit (Southern Biotech).

A stable hTERT RPE-1 cell line expressing CENPA::sfGFP, mCherry::alpha tubulin, and emiRFP670::hCentrin2 was generated using a lentiviral system (Effectene Transfection Reagent, Qiagen) and selected using puromycin, blasticidin, and neomycin. Cells were further selected using fluorescence-activated cell sorting to eliminate CENPA overexpression.

### Mad2 siRNA

Cells were grown to ∼50% confluence in a 10cm dish in DMEM GlutaMAX supplemented with FBS and penicillin-streptomycin (P/S) as described above. 8uL of Lipofectamine RNAiMAX (Thermo Fisher) was added to one tube of 1.2mL of Opti-MEM (Thermo Fisher) and 240pmol of Mad2L1 siRNA (Silencer Select, #4392420) were added to a second tube of 1.2mL OPTIMEM. After 5 minutes of incubation at room temperature with light flicking, the two tubes were combined into one and left to incubate at room temperature for 30 minutes with light flicking. The cells in the 10cm dish were washed once with warm PBS and replaced with 8mL OPTIMEM with 10% FBS. The siRNA mixture (2.4mL total) was added dropwise to the cells. After 24 hours, the cells were washed twice with warm PBS and split onto 25mm glass #1 coverslips in 35mm dishes with DMEM, 10% FBS, and P/S as described above. Cells were imaged the following day, at least 46 hours after siRNA addition.

### Live-cell imaging

All live-cell spinning-disk confocal microscopy imaging was performed as follows. Cells were plated on 25 mm diameter, #1-thickness, round coverglasses (Bioscience Tools) in 35mm dishes to 70-80% confluency (control and UMK57) or 30-40% confluency (monastrol washout) the day before experiments.

For control and UMK57 samples, media was replaced with DMEM containing 1:4000 SPY650-DNA (Cytoskeleton) at least one hour before imaging.

For the monastrol washout samples, mitotic cells were shaken off of the coverslips and the DMEM was replaced with 100µM monastrol (SelleckChem) DMEM. The samples were imaged after 2 hours of incubation in drug media.

Imaging experiments were performed on a home-built spinning disk confocal microscope (Nikon Ti2000, Yokugawa CSU-X1) with 488nm, 561nm, and 647nm lasers, a CMOS camera (Hamamatsu) and a 60x oil immersion objective. Imaging was controlled using a custom MicroManager program. The samples were transferred to a custom-built cell-heater calibrated to 37°C (Bioscience Tools).

For control samples, cells were covered with 750µL of Fluorobrite DMEM (Thermo Fisher) supplemented with 10mM HEPES (Thermo Fisher) and 1µM UMK57 (if relevant; ChemFarm) and covered with 2mL of mineral oil. Up to 22 cells with condensed chromosomes (selected based on SPY650-DNA signal) were imaged in timelapses. Three separate fluorescence channels were acquired every 1 minute with 50ms exposure for 488nm, 546nm, and 647nm excitation in 3 z planes with 3µm spacing. End of nuclear envelope breakdown time was determined based on movies – for cells that underwent nuclear envelope breakdown before the start of imaging, NEBD end times were backtracked using spindle morphologies.

For monastrol washout samples, cells were covered with 750µL of Fluorobrite DMEM supplemented with 10 mM HEPES and 100µM monastrol. 30-45 mitotic cell positions were imaged in timelapses. Two separate fluorescence channels were acquired every 1 minute with 50ms exposure for both 488nm and 546nm excitation in 3 z planes. Monastrol was washed out after 2 minutes of imaging (2×1mL washes) and replaced with 750µl imaging media containing 0.5% v/v DMSO and covered with mineral oil.

For forcing anaphase, 750µL of 50µM AZ-3146 (SelleckChem) imaging media (with or without 1µM UMK57) was added to the samples at the indicated times (final concentration 25µM AZ-3146). Anaphase times were recorded based on the time when kinetochores began separating in the timelapse movies. After all cells divided, high-resolution images were taken of the divided kinetochores for control and UMK57 samples, and both kinetochores and poles for monastrol washout samples (50ms exposure at full laser power for both 488nm and 647nm, 0.5µm z spacing) for at least 3 timepoints, spaced 10 minutes apart.

### Quantitative analysis of kinetochore count data

Z-stacks were run through the custom Python 3 kinetochore counting code in JupyterLab. First, the sfGFP::CENPA channel was masked to separate the cytoplasm from the background, then further masked to separate out the kinetochore pixels from the cytoplasm. Each kinetochore pixel “walked” to its brightest neighboring pixel until the positions converged. Out of the final positions, pixels that had fewer than 100 walkers that ended up at the position and a pixel intensity below 0.0005 were filtered out, leaving candidate kinetochore centers. 3D Gaussian fits were performed on each candidate kinetochore center, and a histogram of integrated intensities were plotted. Final kinetochore count numbers were determined by manual investigation of outliers in 3D Gaussian fitting. Final kinetochore count numbers for each cell were recorded in a spreadsheet and imported into JupyterLab for further analysis. Kinetochore counting code can be found on Github (https://github.com/gloriaha/kinetocounter).

### Fitting of forced anaphase time course data

For fitting the control and UMK57 samples, cells were binned into two minute intervals, and means and standard deviations of the squared kinetochore count differences were calculated for each bin (Figures 2F, 4D). We chose two minute bins due to the 1 minute intervals between subsequent images of a single cell, and the addition of Mps1 inhibitor in between subsequent frames. Assuming statistical symmetry in initial erroneous attachments (i.e. that initially erroneous attachments are equally likely to be associated with either pole) leads to the following prediction for the dynamics of ⟨(Δ*N*)^2^⟩ (SI Appendix):

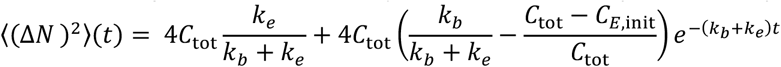

where *t* is time in minutes between NEBD end and Mps1i addition and ⟨(Δ*N*)^2^⟩(*t*)is the predicted mean squared kinetochore count difference at time t. The resulting residual expression was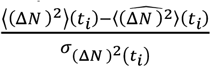, where 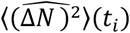 is the empirical mean and 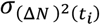 is the empirical standard deviation of (Δ*N*)^2^ at time *t*_i_. We used Scipy’s curve_fit function on the summed squared residuals for each bin to fit the parameter combinations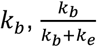, and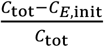. We used the resulting covariance matrix to estimate the propagated parameter fit errors and 2 σ confidence interval (shaded region of plot) using Numpy’s uncertainty module.

For the monastrol samples, cells were divided into timepoints based on the time elapsed between monastrol washout and Mps1i addition, and means and standard deviations of the squared kinetochore count differences were calculated for each bin (Figure 4H). Since the initial ⟨(Δ*N*)^2^⟩ was too large to be consistent with statistical symmetry in initial erroneous attachments, we used a model which account for initial statistical asymmetry (i.e. that cells can contain a “stronger” pole that initially erroneous attachments are more likely to be associated with), leading to the following prediction for the dynamics of ⟨(Δ*N*)^2^⟩ (SI Appendix), which we used to fit the data:

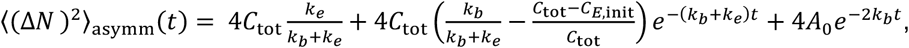

where *A*_0_ is a term that encapsulates asymmetry in the initial erroneous attachments (see SI Appendix).

The resulting residual was 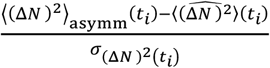, which was then used in the simultaneous fits (see below). For the individual forced anaphase data fit for the monastrol data, we used the approximation

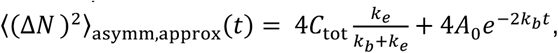

since the high initial ⟨(Δ*N*)^2^⟩ values indicate that this second exponential term strongly dominates under these conditions.

### Fitting of anaphase onset time data

Timelapse movies with 1 minute resolution were used to estimate the anaphase onset time relative to NEBD end (Figure 3B). The first frame where the sister kinetochores started moving apart was taken to be the anaphase onset time. Data was fit to the predicted distribution of anaphase times (SI Appendix):

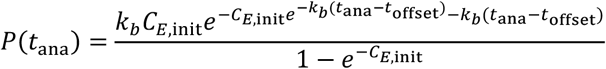

Since cells were imaged in 1 minute intervals, anaphase onset time data was divided into 1 minute bins and each bin was normalized by the total number of cells. The resulting residual expression was *y*_hist,i_ – *P*(*x*_hist,i_), where *y*_hist,i_ was the fraction of cells in the bin starting at time *x*_hist,i_. The best fit *k*_*b*_, *C*_*E*,init_, and *t*_offset_ were determined using Scipy’s curve_fit function on the summed squared residuals for each bin and the resulting Jacobian was used to estimate the parameter fit errors.

### Simultaneous fitting of time course and anaphase onset data (statistically symmetric initially erroneous attachments and neglecting the contribution of finite error rate for anaphase times)

The simultaneous fits for control and UMK57 data in Fig 4D (Table S1) were done using a joint squared residual expression, where data points were normalized by the number of time points (exponential) or number of anaphase time data points (Gumbel). We chose to normalize the data in this way in order to equally weight the two datasets (forced anaphase kinetochore counts and spontaneous anaphase times) when fitting for the maximally likely parameters that fit both datasets.

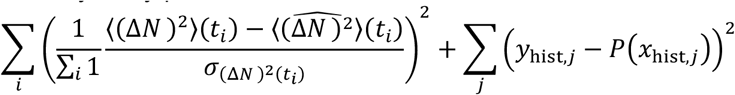

The best fit *k*_*b*_, *k*_*e*_, *C*_*E*,init_, *t*_offset_ and *A*_0_ were determined using Scipy’s curve_fit function on the summed squared residuals. We used the resulting covariance matrix to estimate the parameter fit errors and 2*σ* confidence interval (shaded region of plot) using Numpy’s uncertainty module.

### Simultaneous fitting of time course and anaphase onset data (statistically asymmetric initially erroneous attachments and accounting for the contribution of finite error rate for anaphase times)

The simultaneous fit for monastrol washout data (Fig 4H, Table S1) was done using a joint squared residual expression as above, with the expression valid for statistically asymmetric initially erroneous attachments and the numerically simulated probability distribution of the slowest first passage time with a finite error rate (as described in SI Appendix). Performing fits of the control and UMK57 data allowing for statistically asymmetric initially erroneous attachments and accounting for the contribution of finite error rate for anaphase times produced similar results as the case with statistical symmetry and neglecting finite error rate (Table S1).

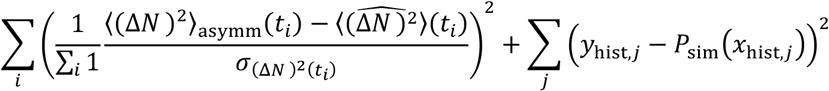

The best fit *k*_*b*_, *k*_*e*_, *C*_*E*,init_, *t*_offset_, and *A*_0_ were determined using Scipy’s curve_fit function on the summed squared residuals. We used the resulting covariance matrix to estimate the parameter fit errors and 2 σ confidence interval (shaded region of plot) using Numpy’s uncertainty module.

## Acknowledgments

We thank Timothy Mitchison, Iain Cheeseman, Vinothan Manoharan, Alexey Khodjakov, Andrea Musacchio, Luyi Qiu, William Conway, Maya Anjur-Dietrich, Easun Arunachalam, and Mustafa Basaran for helpful discussions, and the Bauer Flow Cytometry Core for cell sorting. G.H. was supported by the QuantBio Student Fellowship through the NSF-Simons Center for Mathematical and Statistical Analysis of Biology at Harvard (#1764269). This work was funded by NSF award DBI-1919834. Research reported in this publication was supported by an award from the Zuckerman Travel and Research STEM Fund at Harvard University. A.A. thanks support from the Clore Center for Biological Physics. We also acknowledge support from the CCBX program of the Center for Computational Biology of the Flatiron Institute.

## Author Contributions

G.H. and D.J.N. designed research; G.H. and P.D. performed research; P.D., H.S., and A.A. contributed new analytic tools; G.H. and P.D. analyzed data; and G.H., P.D., A.A., and D.J.N. wrote the paper.

