## Supplementary material for "Measuring and modeling the dynamics of mitotic error correction": SI Appendix

### Supporting Information Text

### Calculation of error correction dynamics in the coarse-grained model

Consider a single chromosome that can either be correctly attached to both poles (bioriented, $B$), erroneously attached to the pole 1 ($E_1$) or erroneously attached to pole 2 ($E_2$). The $E_1$ and $E_2$ states transition to the $B$ state with rate $k_b$ while the $B$ state transitions to the $E_1$ and $E_2$ states with rate $k_e/2$. The probability for a single chromosome to be in states $B$, $E_1$, or $E_2$ evolves according to

$$\dot{\vec{p}} = \dot{\begin{pmatrix} p_{E_1} \\ p_{E_2} \\ p_B \end{pmatrix}} = \begin{pmatrix} -k_b & 0 & k_e/2 \\ 0 & -k_b & k_e/2 \\ k_b & k_b & -k_e \end{pmatrix} \vec{p} = \mathbf{M}\vec{p} \Longrightarrow \vec{p} = e^{\mathbf{M}t}\vec{p_0}, \qquad [1]$$

with $\vec{p_0}$ the initial condition.

Calculating $e^{\mathbf{M}t}$ reveals (with $A \equiv \frac{k_b}{k_b+k_e}$ and $r \equiv k_b + k_e$)

$$e^{\mathbf{M}t} = \frac{1}{2}\begin{pmatrix} 1-A & 1-A & 1-A \\ 1-A & 1-A & 1-A \\ 2A & 2A & 2A \end{pmatrix} + e^{-k_b t}\begin{pmatrix} 1 & -1 & 0 \\ -1 & 1 & 0 \\ 0 & 0 & 0 \end{pmatrix} + e^{-rt}\begin{pmatrix} A & A & A-1 \\ A & A & A-1 \\ -2A & -2A & 2(1-A) \end{pmatrix}. \qquad [2]$$

Thus the steady state is given by

$$\vec{p_{ss}} = \frac{1}{2}\begin{pmatrix} 1-A \\ 1-A \\ 2A \end{pmatrix} \qquad [3]$$

and the temporal dynamics with initial condition $\vec{p_0} = (a, b, c)^T$ as

$$\vec{p}(t) = \vec{p_{ss}} + \frac{1}{2}e^{-k_b t}\begin{pmatrix} a-b \\ b-a \\ 0 \end{pmatrix} + \frac{1}{2}e^{-rt}\begin{pmatrix} A-c \\ A-c \\ 2c-2A \end{pmatrix} \qquad [4]$$

In order to use this framework to model a population of cells, each with many chromosomes acting independently, we construct a random variable called $X = 2(\mathbf{1}_{E_1} - \mathbf{1}_{E_2})$ (where $\mathbf{1}_{E_k}$ is an indicator random variable, taking the value 1 if the chromatid ended in state $E_k$ and 0 otherwise). $X$ is the difference between the number of chromatids that end up in either daughter cell. If both sister chromatids of a chromosome go to daughter cell 1, $X$ is 2, if both go to daughter cell 2, $X$ is -2, and if they split evenly, $X$ is 0. Thus the squared chromatid count difference for a single cell is given by

$$(\Delta N)^2(t) = (\Sigma_i X_i)^2 \qquad [5]$$

which has an expected value of

$$\langle(\Delta N)^2(t)\rangle = \left\langle(\Sigma_i X_i)^2\right\rangle = \Sigma_i\langle X_i^2\rangle + 2\Sigma_i\Sigma_{j>i}\langle X_i X_j\rangle, \qquad [6]$$

the mean squared difference in chromatid (or kinetochore) counts between daughter cells.

Since $X_i X_j$ is 4 with probability $p_{E_2,i}p_{E_2,j} + p_{E_1,i}p_{E_1,j}$ and -4 with probability $p_{E_2,i}p_{E_1,j} + p_{E_1,i}p_{E_2,j}$, the expected value for $i \neq j$ evalutes to

$$\langle X_i X_j\rangle = 4(p_{E_2,i} - p_{E_1,i})(p_{E_2,i} - p_{E_1,i}) = 4e^{-2k_b t}(a_i - b_i)(a_j - b_j). \qquad [7]$$

For $i = j$, $X_i^2$ is 4 with probability $p_{E_2,i} + p_{E_1,i}$ and 0 otherwise, giving

$$\langle X_i^2\rangle = 4(p_{E_2,i} + p_{E_1,i}) = 4[1 - A + e^{-rt}(A - c_i)] \qquad [8]$$

Combining these two results yields the full expression for $\langle(\Delta N)^2(t)\rangle$:

$$\begin{aligned} \langle(\Delta N)^2(t)\rangle &= 4C_{\text{tot}}(1-A) + 4e^{-rt}\left(C_{\text{tot}}A - \Sigma_i c_i\right) + 8e^{-2k_b t}\Sigma_{i,j>i}(a_i - b_i)(a_j - b_j) \\ &= 4C_{\text{tot}}\frac{k_e}{k_b+k_e} + 4C_{\text{tot}}e^{-(k_b+k_e)t}\left(\frac{k_b}{k_b+k_e} - \langle c_i\rangle_c\right) + 4e^{-2k_b t}\left(C_{\text{tot}}^2\langle a_i - b_i\rangle_c^2 - C_{\text{tot}}\langle(a_i - b_i)^2\rangle_c\right), \end{aligned} \qquad [9]$$

where in the last step we used the following relations:

$$C_{\text{tot}}A - \Sigma_i c_i = N\left(A - \frac{1}{C_{\text{tot}}}\Sigma_i c_i\right) = C_{\text{tot}}(A - \langle c_i\rangle_c) \qquad [10]$$

and

$$2\Sigma_{i,j>i}(a_i - b_i)(a_j - b_j) = [\Sigma_i(a_i - b_i)]^2 - \Sigma_i(a_i - b_i)^2 = C_{\text{tot}}^2\langle a_i - b_i\rangle_c^2 - C_{\text{tot}}\langle(a_i - b_i)^2\rangle_c \qquad [11]$$

where $\langle . \rangle_c$ denotes the average over chromosomes in a given cell.

Since we are dealing with an inhomogeneous population of cells (and thus $a_i, b_i$ are not necessarily the same for all cells), we can replace the parameters that depend on the initial condition with population averages:

$$\langle(\Delta N)^2(t)\rangle = 4C_{\text{tot}}\frac{k_e}{k_b+k_e} + 4C_{\text{tot}}e^{-(k_b+k_e)t}\left(\frac{k_b}{k_b+k_e} - \overline{\langle c_i\rangle_c}\right) + 4e^{-2k_b t}\left(C_{\text{tot}}^2\overline{\langle a_i - b_i\rangle_c^2} - C_{\text{tot}}\overline{\langle(a_i - b_i)^2\rangle_c}\right), \qquad [12]$$

We define $C_{E,\text{init}}$ as the population average number of initially erroneous chromosomes per cell, so $\overline{\langle c_i \rangle_c} = \frac{C_{\text{tot}} - C_{E,\text{init}}}{C_{\text{tot}}}$. The amplitude of the $-2k_b$ exponential in the equation for $\langle(\Delta N)^2(t)\rangle$ depends on the degree of statistical symmetry in initial erroneous attachment. In other words, if the initial probability of being erroneously attached to one pole vs. the other is equivalent, then $a_i - b_i$ would be close to 0 for all chromosomes, and thus the magnitude of the $2k_b$ exponential would be close to 0 (statistical symmetry). If instead some chromosomes are more likely to be initially erroneously attached to one pole over the other, the magnitude of the $2k_b$ exponential would not be negligible (statistical asymmetry). If we assume symmetry in initial erroneous attachments on average, we recover the single exponential equation that we use to fit the control and UMK57 data in the main text:

$$\langle(\Delta N)^2(t)\rangle = 4C_{\text{tot}}\frac{k_e}{k_b+k_e} + 4C_{\text{tot}}e^{-(k_b+k_e)t}\left(\frac{k_b}{k_b+k_e} - \frac{C_{\text{tot}} - C_{E,\text{init}}}{C_{\text{tot}}}\right). \quad [13]$$

For $C_{\text{tot}} = 46$ chromosomes, the maximum possible value of $\langle(\Delta N)^2(t=0)\rangle$ assuming statistical symmetry in initial erroneous attachments is $4C_{\text{tot}} = 184$ (this is the case when $a = b = 1/2$ and $c = 0$, corresponding to binomial partitioning of the chromatids). Given that $\langle(\Delta N)^2(t=5)\rangle$ for the monastrol washout is 558, we know that $4C_{\text{tot}}^2\langle a_i - b_i\rangle_c^2$ must be non-zero, indicating that there must be a left-right symmetry breaking on average. Thus we use the full double exponential equation to fit the monastrol washout data, where we lump the asymmetry coefficient into one constant $A_0$:

$$\langle(\Delta N)^2(t)\rangle = 4C_{\text{tot}}\frac{k_e}{k_b+k_e} + 4C_{\text{tot}}e^{-(k_b+k_e)t}\left(\frac{k_b}{k_b+k_e} - C_{E,\text{init}}\right) + 4A_0e^{-2k_bt}, \quad [14]$$

Fitting the control and UMK57 to this full model results in a best fit with with $A_0$ indistinguishable from zero (Table S1), indicating that there is no discernable initial statistical asymmetry in these samples. This is consistent with the observation that most RPE-1 cells begin with centrosomes (spindle "poles") aligned on the short nuclear axis, equidistant from a nuclear envelope packed with chromosomes (1). In such an initial configuration, one wouldn't expect that one pole would be preferred for chromosome attachments over the other. On the other hand, in the monastrol initial condition, the centrosomes are positioned in the center of a ring of chromosomes, and most chromosomes are only attached to one of the centrosomes (2), which could lead to initial statistical asymmetry. Differences in factors such as the movement of poles in opposite directions during spindle bipolarization could also lead to one pole having more erroneous attachments than the other.

### Calculation of anaphase onset times in the coarse-grained mode

We used the same coarse-grained model described above to predict the distribution of anaphase onset times. If transitions from the correct state (state $B$) to the erroneous states (the $E$ states) can be neglected (i.e. if $k_e \approx 0$), then the distributions of anaphase times can be calculated analytically. Alternatively, if it is necessary to account for finite $k_e$, then the distributions of anaphase times must be calculated numerically. We treat these two cases in turn below.

**Anaphase onset time distribution for $k_e \approx 0$.** When $k_e = 0$, then if a chromosome is incorrect at time $t = 0$, its probability of remaining incorrect at time $t$ decays exponentially, and the probability of being correct is:

$$C(t) = 1 - e^{-k_bt}. \quad [15]$$

If there are $C_{E,\text{init}}$ initially erroneous chromosomes, then the probability that *all* of them will be corrected by time $t$ is therefore:

$$G(t) = (1 - e^{-k_bt})^{C_{E,\text{init}}} \approx e^{-C_{E,\text{init}}e^{-k_bt}}, \quad [16]$$

(see chapter 6 in Ref. (3) for a discussion on the accuracy of this approximation).

The probability distribution (also referred to as the probability density function), is then given by the derivative of $G(t)$, yielding a Gumbel distribution:

$$p(t_f) \approx k_bC_{E,\text{init}}e^{-C_{E,\text{init}}e^{-k_bt_f} - k_bt_f}. \quad [17]$$

If we assume that the anaphase time is given by the slowest first passage time plus the offset time ($t_{\text{ana}} = t_f + t_{\text{offset}}$), we can finally express the distribution of anaphase onset times as

$$p_{C_{E,\text{init}}}(t_{\text{ana}}) = k_bC_{E,\text{init}}e^{-C_{E,\text{init}}e^{-k_b(t_{\text{ana}}-t_{\text{offset}})} - k_b(t_{\text{ana}}-t_{\text{offset}})}. \quad [18]$$

Thus, if $k_e \approx 0$, then this coarse-grained model predicts that the distribution of anaphase times should follow a (shifted) Gumbel distribution. Since our measurements show that the rate at which erroneous attachments are corrected is much faster that the rate at which correct attachments become erroneous, it is reasonable to expect that this will be a good approximation.

To find the expectation value (the mean of the last passage time), we can integrate this probability density.

$$E[t_f] = \int_{-\infty}^{\infty} t_f(k_bC_{E,\text{init}}e^{-C_{E,\text{init}}e^{-k_bt_f} - k_bt_f})\mathrm{d}t_f = \frac{\gamma + \log(C_{E,\text{init}})}{k_b} \quad [19]$$

$\gamma \approx 0.5772$ is Euler's constant, yielding the following approximate equation for the mean anaphase time:

$$E[t_{\text{ana}}] = E[t_f] + t_{\text{offset}} = \frac{\gamma + \log(C_{E,\text{init}})}{k_b} + t_{\text{offset}} \quad [20]$$

Thus the mean anaphase time depends on the correction rate $k_b$, and can be shifted by $t_{\text{offset}}$ and $C_{E,\text{init}}$.

We can calculate the variance similarly. First, we take the expectation value of the squared slowest first passage time.

$$\int_{-\infty}^{\infty} t_f^2 (k_b C_{E,\text{init}} e^{-C_{E,\text{init}} e^{-k_b t_f} - k_b t_f}) \mathrm{d}t_f = \frac{6\gamma^2 + \pi^2 + 6\log(C_{E,\text{init}})(2\gamma + \log(C_{E,\text{init}}))}{6k_b^2} \quad [21]$$

Then we calculate the variance:

$$\begin{aligned} \text{Var}[t_{\text{ana}}] = \text{Var}[t_f] = E[t_f^2] - E[t_f]^2 &= \frac{6\gamma^2 + \pi^2 + 6\log(C_{E,\text{init}})(2\gamma + \log(C_{E,\text{init}}))}{6k_b^2} - \left(\frac{\gamma + \log(C_{E,\text{init}})}{k_b}\right)^2 \\ &= \frac{\pi^2}{6k_b^2} \end{aligned} \quad [22]$$

The variance is completely independent of $C_{E,\text{init}}$ and $t_{\text{offset}}$, while it scales with $1/k_b^2$. Thus the width of the Gumbel distribution is entirely determined by the error correction rate $k_b$, so we can constrain $k_b$ by fitting the observed anaphase times to the Gumbel distribution.

**Anaphase onset time distribution for finite $k_e$.** If $k_e$ is finite, and cannot be well approximated as zero, then the distribution of anaphase time needs to be solved numerically. To do so, we first defined an initial condition vector $\vec{v}$ of length $C_{\text{tot}}$ with integer states going from $i = 0$ (all chromosomes incorrectly attached in state E) to $i = C_{\text{tot}} - 1$ (all but one chromosome in state B). $\vec{v}$ was defined as a binomial distribution with 46 trials and probability $\frac{C_{\text{tot}} - C_{E,\text{init}}}{C_{\text{tot}}}$. Thus for a given $c = \frac{C_{\text{tot}} - C_{E,\text{init}}}{C_{\text{tot}}}$, or initial probability of a chromosome being correctly attached, $\vec{v}$ represents the probability distribution of the number of correctly-attached chromosomes.

We then defined transition rates between states as follows:

$$\begin{aligned} i, i+1 &: k_b(C_{\text{tot}} - i) \\ i, i-1 &: k_e i \\ i, i &: 1 - k_b(C_{\text{tot}} - i) - k_e i. \end{aligned}$$

We used these transition rates to populate a $(C_{\text{tot}} \times C_{\text{tot}})$ transition matrix $\mathbf{A}$. We then calculated the probability of having corrected all attachments by a given time $t$ as

$$P(t_f \leq t_i) = 1 - \sum e^{\mathbf{A} t_i} \vec{v}, \quad [23]$$

and thus the probability of the slowest first passage time being in a time interval $\{t_i, t_{i-1}\}$ was given by:

$$P(t_{i-1} < t_f \leq t_i) = \frac{P(t_f \leq t_i) - P(t_f \leq t_{i-1})}{t_i - t_{i-1}}. \quad [24]$$

Finally, we calculated the probability of the anaphase onset time being in a given time interval as:

$$P(t_{i-1} + t_{\text{offset}} < t_{\text{ana}} \leq t_i + t_{\text{offset}}) = \frac{P(t_{\text{ana}} \leq t_i + t_{\text{offset}}) - P(t_{\text{ana}} \leq t_{i-1} + t_{\text{offset}})}{t_i - t_{i-1}}. \quad [25]$$

For a given set of parameters $k_e$, $k_b$, $C_{E,\text{init}}$, and $t_{\text{offset}}$, we calculated the numerical probability distribution of anaphase onset times and compared it to the data in order to solve for the parameters that best fit the data.

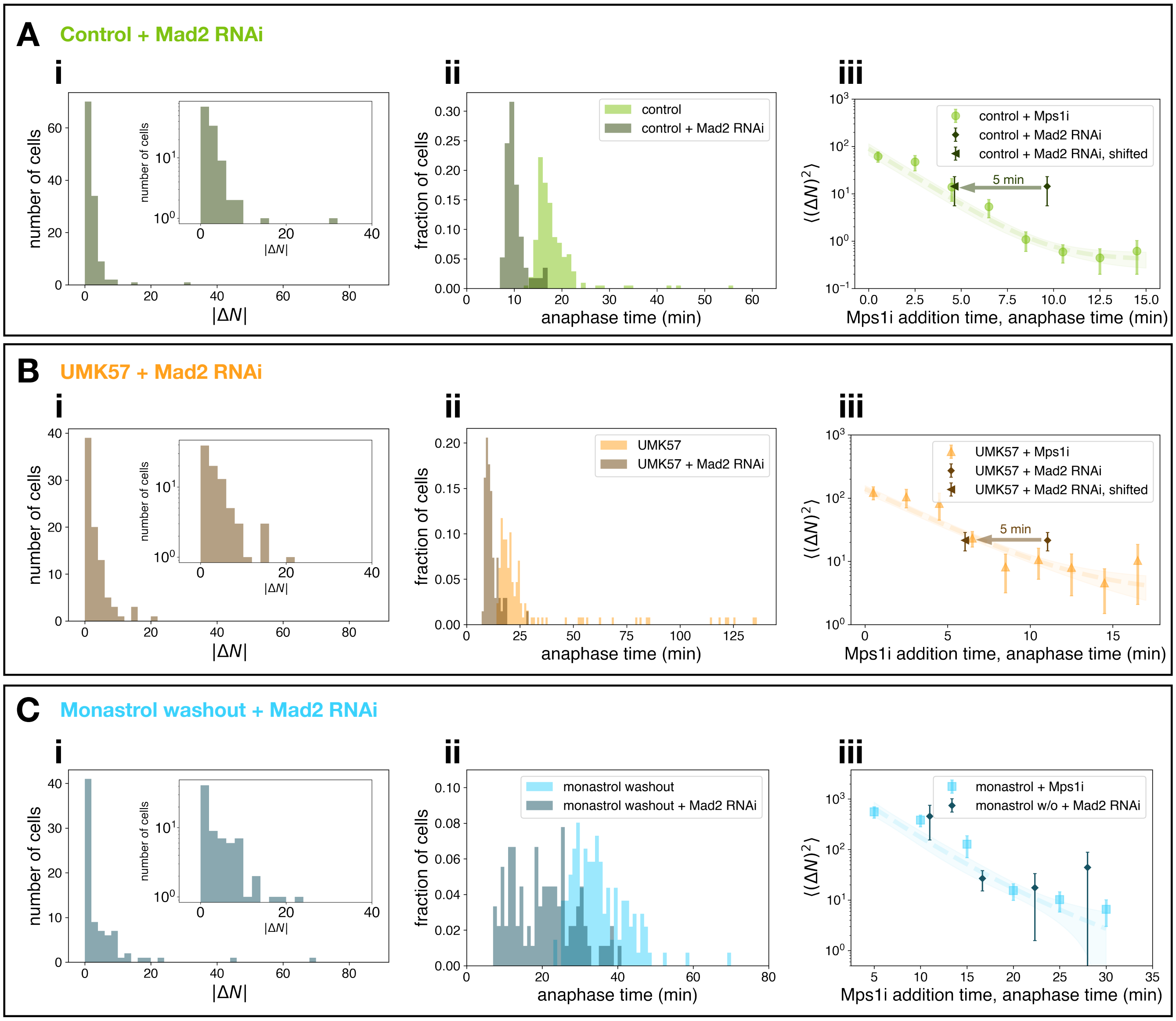


**Fig. S1.** Mad2 RNAi results. (A) (i) Unforced (no Mps1i added) kinetochore count difference distribution for control RPE1 cells with Mad2 RNAi knockdown (n=119, inset: zoomed in with logged y axis). (ii) Unforced anaphase times for control RPE1 cells with (n=57) and without Mad2 RNAi knockdown. (iii) Mean squared kinetochore count difference vs. mean anaphase onset time for unforced control cells with Mad2 RNAi knockdown overlaid on forced anaphase $\langle(\Delta N)^2\rangle$ curve for control cells without Mad2 RNAi. Shifted point is shifted 5 minutes before the mean anaphase time. (B) Unforced $|\Delta N|$ for Mad2 RNAi cells with 1uM UMK57 (n=85), unforced anaphase times for 1uM UMK57 cells with (n=68) and without Mad1 RNAi, $\langle(\Delta N)^2\rangle$ vs. mean anaphase time for Mad2 RNAi+UMK57 cells overlaid on forced anaphase $\langle(\Delta N)^2\rangle$ curve. Shifted point in (iii) is shifted 5 minutes before the mean anaphase time for UMK57 Mad2 RNAi cells. (C) Unforced $|\Delta N|$ for Mad2 RNAi cells after washout from monastrol incubation (n=78), anaphase onset times relative to monastrol washout times for cells with (n=90) and without Mad2 RNAi, $\langle(\Delta N)^2\rangle$ vs. mean anaphase time for Mad2 RNAi monastrol washout cells binned in 5 minute bins overlaid on forced anaphase $\langle(\Delta N)^2\rangle$ curve for monastrol washout cells, all times measured relative to monastrol washout time. Error bars are standard error of the mean.

**Table S1. Simultaneous fit results**

| Condition | Model (initial attachments, anaphase time) | $k_b$ (min$^{-1}$) | $k_e$ (min$^{-1}$) | $C_{E,\mathrm{init}}$ | $t_{\mathrm{offset}}$(min) | $A_0$ |
|---|---|---|---|---|---|---|
| **control** | symmetric, $k_e \approx 0$ | $0.55 \pm 0.02$ | $0.001 \pm 0.0002$ | $26 \pm 3$ | $10.5 \pm 0.3$ | N/A |
| | asymmetric, finite $k_e$ | $0.57 \pm 0.03$ | $0.001 \pm 0.0003$ | $26 \pm 9$ | $10.8 \pm 0.5$ | $0 \pm 13$ |
| **UMK57** | symmetric, $k_e \approx 0$ | $0.28 \pm 0.01$ | $0.005 \pm 0.002$ | $35 \pm 2$ | $5.2 \pm 0.5$ | N/A |
| | asymmetric, finite $k_e$ | $0.34 \pm 0.03$ | $0.005 \pm 0.002$ | $41 \pm 3$ | $6.5 \pm 0.6$ | $0 \pm 16$ |
| **monastrol w/o** | asymmetric, finite $k_e$ | $0.15 \pm 0.03$ | $0 \pm 0.001$ | $46 \pm 51$ | $6 \pm 4$ | $620 \pm 170$ |

 Gloria Ha, Paul Dieterle, Hao Shen, Ariel Amir, and Daniel J. Needleman
